# Compaction of Duplex Nucleic Acids upon Native Electrospray Mass Spectrometry

**DOI:** 10.1101/105049

**Authors:** Massimiliano Porrini, Frédéric Rosu, Clémence Rabin, Leonardo Darré, Hansel Gómez, Modesto Orozco, Valérie Gabelica

## Abstract

Native mass spectrometry coupled to ion mobility spectrometry is a promising tool for structural biology. Intact complexes can be transferred to the mass spectrometer and, if native conformations survive, collision cross sections give precious information on the structure of each species in solution. Based on several successful reports for proteins and their complexes, the conformation survival becomes more and more taken for granted. Here we report on the fate of nucleic acids conformation in the gas phase. Disturbingly, we found that DNA and RNA duplexes, at the electrospray charge states naturally obtained from native solution conditions (≥ 100 mM aqueous NH_4_OAc), are significantly more compact in the gas phase compared to the canonical solution structures. The compaction is observed for short (12-bp) and long (36-bp) duplexes, and for DNA and RNA alike. Molecular modeling (density functional calculations on small helices, semi-empirical calculations on up to 12-bp, and molecular dynamics on up to 36-bp duplexes) demonstrates that the compaction is due to phosphate group self-solvation prevailing over Coulomb-driven expansion. Molecular dynamics simulations starting from solution structures do not reproduce the experimental compaction. To be experimentally relevant, molecular dynamics sampling should reflect the progressive structural rearrangements occurring during desolvation. For nucleic acid duplexes, the compaction observed for low charge states results from novel phosphate-phosphate hydrogen bonds formed across both grooves at the very late stages of electrospray.

## INTRODUCTION

Besides genetic information storage, nucleic acids perform pivotal regulatory functions in living organisms.^1^ Conformational changes are key to these regulation mechanisms, whether at the DNA level (e.g., particular structures in gene promoters)^2^ or at the RNA level (e.g., riboswitches).^3^ Electrospray ionization mass spectrometry (ESI-MS) in native conditions helps deciphering the complexation equilibria.^4-7^ Coupled to ion mobility spectrometry (IMS), native mass spectrometry makes it possible to reveal the ensuing conformational responses.^8-10^ But before native ESI-IMS-MS can be applied to study nucleic acids conformations from solution, it is essential to understand to what extent the different types of DNA/RNA secondary structures are preserved, or affected, by the transition from the solution to the gas phase.

Tandem mass spectrometry^11-14^ experiments have shown that the kinetic stability of gas-phase DNA duplexes was correlated with the fraction of guanine-cytosine (GC) base pairs, and with the base pair ordering. This suggested that *Watson-Crick* (*WC*) hydrogen bonding and base stacking were at least partially maintained in the gas phase. Infrared multiphoton dissociation spectroscopy on duplexes and single strands suggested that GC bases are engaged in hydrogen bonds in the gas-phase duplexes, but no such evidence was found for AT base pairs.^15^ Early molecular dynamics (MD) validated this view: although (12-mer)_2_^6-^ and (16-mer)_2_^8-^ duplexes distort in the gas phase, most *WC* H-bonding and stacking interactions are preserved, particularly in GC-rich regions.^16^

The Bowers group later explored the gas-phase structure of DNA duplexes by IMS.^17,18^ By comparing the experimental collision cross section (CCS) of d(GC)_n_ duplexes with those obtained on duplexes relaxed by short (5-ns) MD, they showed that the gas-phase structures resemble an A-helix for the short duplexes (8— 16-mer) and a B-helix for longer ones (>18-mer). It is worth noting that Bowers’ IMS experiments were performed in non-native solution conditions (49:49:2 mixture of H_2_O:MeOH:NH_4_OH) and on high charge states (1 negative charge per 2 base pairs), raising questions on whether this represents what happens in native conditions. Here we show that, at the lower charge states typically obtained from native solution conditions (100—150 mM NH_4_OAc in 100% H_2_O, pH=7), the conformations are significantly more compact than expected for B- or A-helices. We investigated the physical origin of this gas-phase compaction by molecular modeling.

## METHODS

### Electrospray ion mobility spectrometry

DNA and RNA duplexes were prepared by annealing their corresponding single-strands (purchased from Eurogentec, Seraing, Belgium, with RPcartridge-Gold purification) in aqueous 100 mM NH_4_OAc. When sprayed at 10 µM duplex, the major charge states are 4- for the 10-bp, 5-for the 12-bp, 7- for the 24-bp, 8- and 9- for the 36-bp duplexes. Higher charge states were generated by adding 0.2% to 0.75% sulfolane to the solution. ESI-IMSMS experiments were recorded on an Agilent 6560 IMS-Q-TOF, with the drift tube operated in helium (supporting information Section S1). The arrival time distributions were fitted by Gaussian peaks and the CCS values of the center of each peak were determined by the stepped-field method. For visualization, we converted the arrival time distributions into CCS distributions (see supporting information).

### Gas-phase simulations

The starting structures of the duplexes were built with the Nucleic Acid Builder (NAB) software,^19^ both for the A-form (DNA and RNA) and the B-form (DNA). Table 1 lists the main sequences and levels of theory used here. The numbers (33, 66, and 100) in duplex names reflect the GC content (in %) of each 12-bp unit. For 12-d_33_, 12-d_66_, 12-d_100_ (B-helix) and 12-r66 (A-helix), we first carried out MD simulations in water. Then all water molecules and counterions were removed at once, before each 1-µs gas-phase simulation.

**Table 1.**
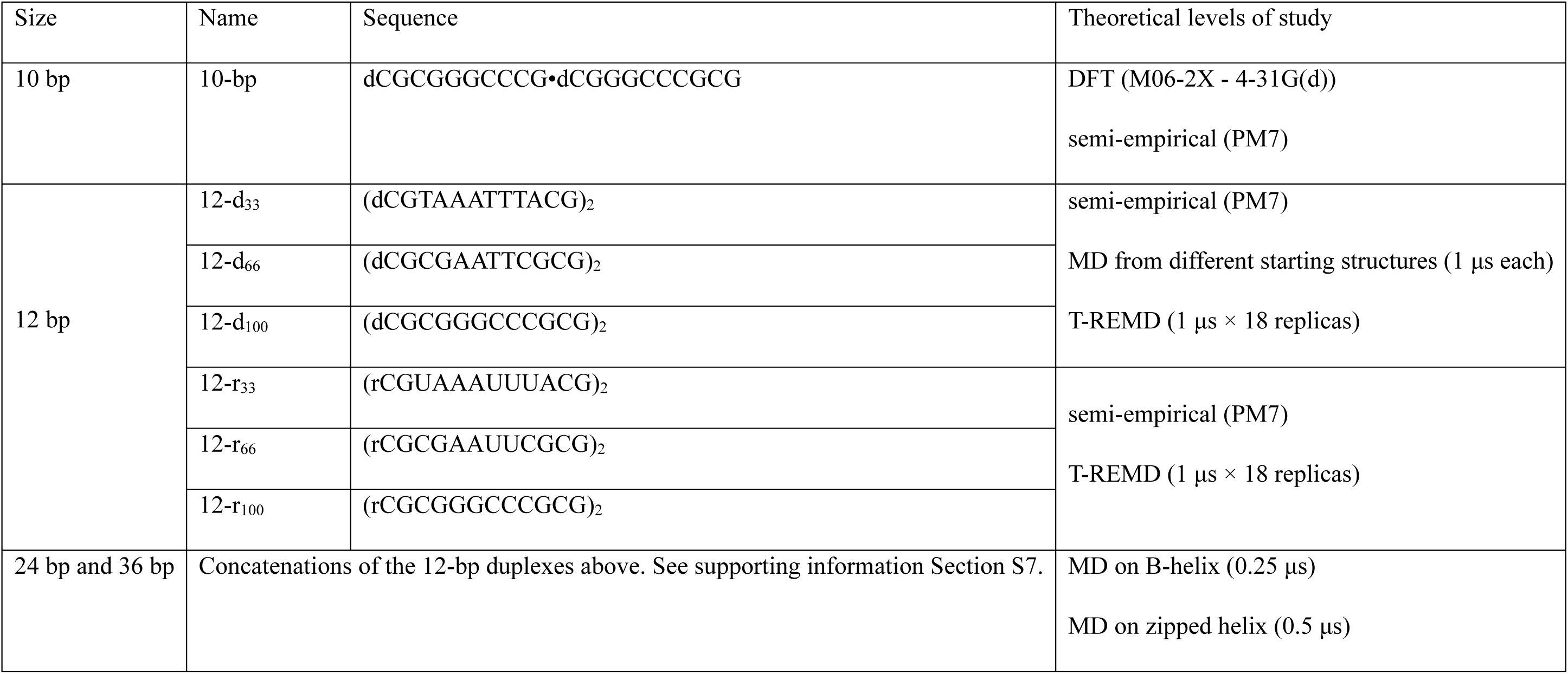
Size, name and sequences of the duplexes under study, and outline of calculations carried out.

The two possibilities to reduce the total charge to −5 (major charge state) are the localized charges (LC) and distributed charges (DC) models.^16^ LC and DC gave similar results upon unbiased MD of duplexes,^16^ so here only the LC model was tested extensively. With the LC model, protons are added on 17 out of the 22 phosphate groups. Among the 26334 possible protonation schemes, we selected a few low-energy ones based on single point molecular mechanics calculations. With the DC model, the net charge of each phosphate group is reduced so that the total charge of the duplex is -5. Because temperature replica exchange MD (T-REMD) simulations on gas-phase duplexes had never been attempted before, LC and DC models were both tested for T-REMD.

Solution and gas phase MD simulations were carried out with the MPI-versions of modules *pmemd* and *sander*, respectively, of the Amber12 suite of programs,^20^ implementing parmBSC1 force field^21^ for DNA and parmBSC0 force field + χ_OL3_ correction^22,23^ for RNA. The electrostatic interactions were calculated with the particle mesh Ewald algorithm^24^ (real-space cut-off = 10 Å) in solution and direct Coulomb summation (no cut-off) in gas phase. All 12-bp duplexes were subjected gas phase T-REMD^25^ (1 µs × 18 replicas) with temperature values from 300.00 to 633.94 K, chosen with predictor from reference 26 (average successful exchange rate of ca. 30%). Short duplexes (7— 10 bp) were optimized at DFT level,^27^ with the M06-2X^28^ functional including the dispersion correction GD3.^29^ The basis set was 6-31G(d,p) for the 7—9-bp duplexes (see Supporting Information) and 4-31G(d) for the 10-bp duplex. Duplexes up to 12-bp were also studied at the semi-empirical (SE) level with MOPAC,^30^ using different methods31 (further details in Supporting Information). Hydrogen bond and stacking analysis was performed for all simulations as detailed in the Supplementary Information section S2.

### Simulation of desolvation and proton transfer

Starting from equilibrated MD simulations in solution, we cut droplets of ca. 2400 water molecules (radius ~ 25 Å) containing the duplex 12-d_66_, and 16, 17 and 18 NH_4_^+^ cations to give a total net charge of −6, −5 and −4 respectively. The droplets were then subjected to gas-phase MD simulations following Konermann’s protocol.^32^ Briefly, the trajectories were propagated by 500-ps chunks at constant temperature (350 K). To accelerate the evaporation, at the beginning of each chunk the initial velocities were reassigned according to the Boltzmann distribution at T = 350 K. At the end of each chunk, we stripped out all water molecules farther than 60 Å from the N6 atom of the 18^th^ residue adenine (which is approximately in the center of the duplex). A further 50-ns chunk at T = 450 K helped the last “sticky” water molecules to evaporate. In total, twelve independent trajectories were obtained (four at each charge state, 4-, 5- and 6-). We then localized the charges (LC model) as follows: on the ultimate conformation of every trajectory a proton from each NH_4_^+^ cation was transferred to the closest phosphate oxygen atom, and ammonia is removed. The resulting duplexes were then subjected to 1-µs unbiased MD.

### CCS calculations

The collision cross section (CCS) is calculated using the EHSSrot code^33^ with the atom parameterization of Siu et al,^34^ a combination that is both accurate and efficient for calculating the CCS of nucleic acids in the gas phase.^35^ The CCS is calculated for snapshots every 0.5 ns in each MD trajectory.

## RESULTS AND DISCUSSION

### 1) DNA duplexes in their predominant native charge state are more compact than a canonical B-helix.

When sprayed from aqueous 100 mM NH_4_OAc, the major charge state of 12-bp duplexes is 5- (Figure S2). The ^DT^CCS_He_ distributions for the 5- duplexes without ammonium ions bound and in the softest ion transfer conditions are shown in Figure 1A (full results in supporting Figure S3). These distributions suggest more compact structures than those of canonical B- or A-forms (Table 2). Upon pre-IMS activation, the CCS distribution of 12-d_33_ is unchanged, 12-d66 is losing the high-CCS peak to the profit of the low-CCS peak, and the entire distribution of 12-d_100_ is shifting towards lower CCS, down to ~705 Å² (see supporting information Figure S4). At low pre-IMS activation, duplexes^5-^ with ammonium adducts are also detected. The CCS of these ions is similar to that of the bare duplexes, but for 12-d_66_ the higher-CCS peak is more abundant–see supporting Figure S5). The peak center values obtained for soft and harsh conditions are listed in Table 1.

**Figure 1.**
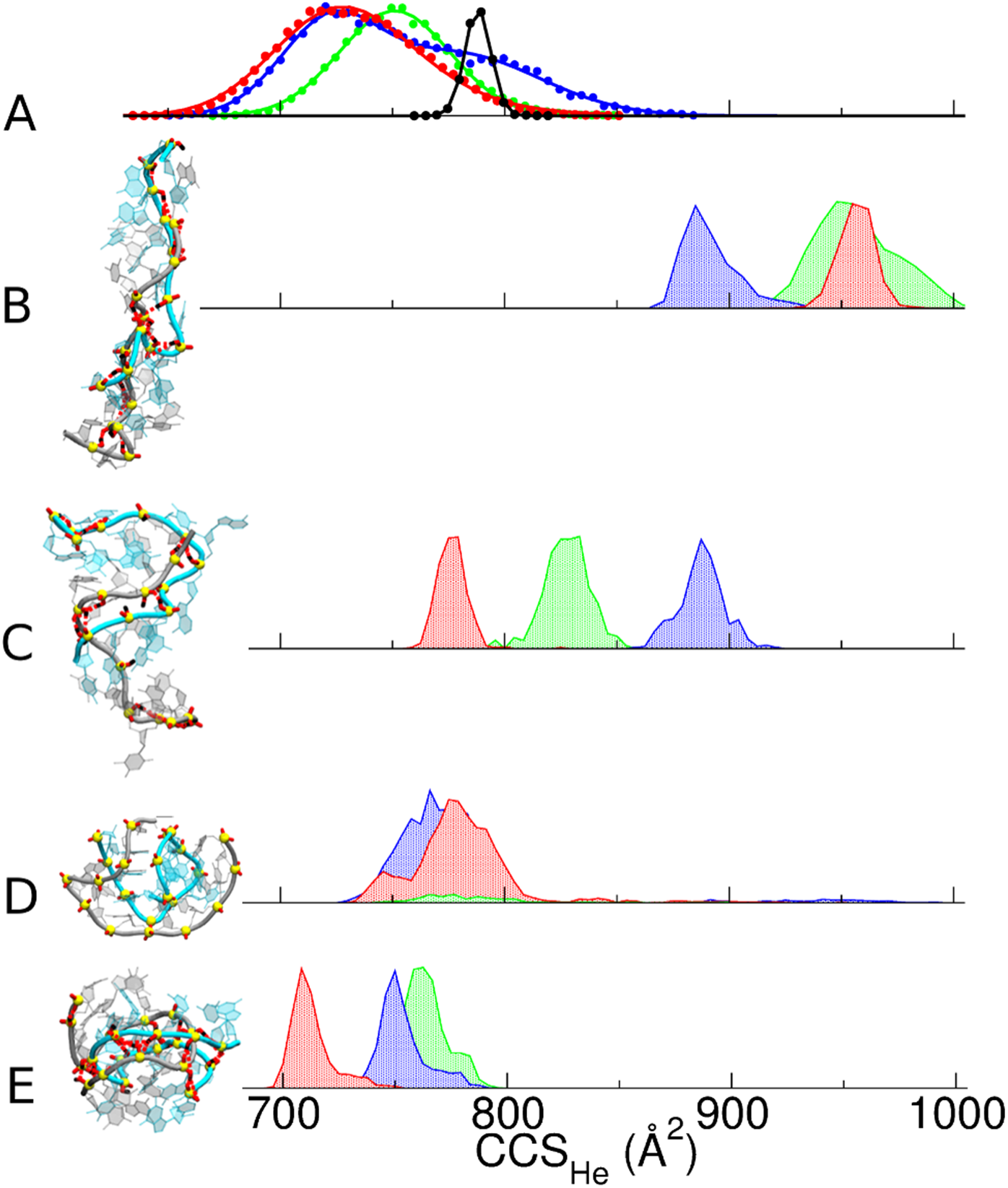
(A) Experimental DTCCS_He_ distribution for the 12-bp DNA duplexes (green: (12-d_33_)^5-^, blue: (12-d_66_)^5-^, red: (12-d_100_)^5-^) and the rigid G-quadruplex ([d(TG_4_T)]_4_)^5-^ (black). (B-E) Calculated CCS distributions for molecular models generated by (B) gas-phase MD of the B-helices, (C) gas-phase MD of the A-helices, (D) T-REMD simulations on B-helices using distributed charges (note that most of the population of (12-d_33_)^5-^ duplex dissociated during simulations), and (E) MD following a restrained minimization forcing H-bond formation between the phosphate groups across both grooves of a B-helix. The final MD structures of each duplex model, created with VMD software,^49^ are shown for (12-d66)^5-^ on the same scale (see supporting Figure S10 for a magnification).

**Table 2.** Collision cross section values in helium (CCS_He_) of 12-bp duplexes, prior and subsequent to PM7 optimization, unbiased MD, TREMD (DC model; results with LC model are not listed because they depend on the chosen locations–see text), and unbiased MD on the helix zipped by restrained minimization. Errors reported for MD are the standard deviation on the different structures along the trajectory.

| | $\text{DTCCS}_{\text{He}} (\text{\AA}^2)$ | $\text{CALCCCS}_{\text{He}} (\text{\AA}^2)$ | | | | | | | | |
| --- | --- | --- | --- | --- | --- | --- | --- | --- | --- | --- |
|  |  | B-helix |  |  |  | A-helix |  |  |  | Zipped helix |
|  |  | Canonical | PM7 | MD | T-REMD (DC) | Canonical | PM7 | MD | T-REMD (DC) | MD |
| $[\text{12-d}_{33}]^{5-}$ | Soft/harsh: 760 | 918 | 914 | $954 \pm 18$ | $780 \pm 24$ | 900 | 861 | $826 \pm 11$ | | $759 \pm 12$ |
| $[\text{12-d}_{66}]^{5-}$ | Soft: 735/803<br>Harsh : 735 | 903 | 831 | $881 \pm 6$ | $770 \pm 15$ | 893 | 942 | $886 \pm 10$ | | $752 \pm 10$ |
| $[\text{12-d}_{100}]^{5-}$ | Soft: 730<br>Harsh: 710 | 908 | 838 | $953 \pm 8$ | $771 \pm 14$ | 892 | 869 | $774 \pm 6$ | | $711 \pm 9$ |
| $[\text{12-r}_{33}]^{5-}$ | - | - | - | | | 944 | 938 | | $741 \pm 21$ | |
| $[\text{12-r}_{66}]^{5-}$ | Soft/harsh: 725 | - | - | | | 945 | 902 | | $747 \pm 18$ | |
| $[\text{12-r}_{100}]^{5-}$ | Soft: 735<br>Harsh: 695 | - | - | | | 940 | 896 | | $743 \pm 11$ | |

Charge states 4- and 6- are also detected in aqueous NH_4_OAc. The duplex CCS distributions are highly charge-state dependent (see supplementary Figure S3): charge states 4- and 5- are similarly compact, whereas charge state 6- has a ~20% larger CCS. Charge state 7-, obtained for 12-d_66_ and 12-d100 by adding the “super-charging” agent sulfolane,^36^ has a >30% larger CCS.

The duplex CCS distributions are significantly broader than those of the tetramolecular G-quadruplex [dTG_4_T]_4_ (in black in Figure 1A, and supplementary Figure S3), a rigid structure of the same size. This indicates that a greater conformational space is explored in the gas phase by nucleic acid duplexes compared to the G-quadruplex,^14^ and that gas-phase duplexes consist of an ensemble of conformations not fully interconverting on the time scale of the mobility separation (10-30 ms).

### 2) MD trajectories, DFT and semi-empirical optimizations reveal phosphate-phosphate hydrogen bond formation.

To find out which tridimensional structures are compatible with the experimental CCS values of the 12-bp^5-^ duplexes, we first carried out unbiased MD simulations directly from B- and A-helix structures, stripped of the solvent. Figure 1B,C shows representative CCS distributions. All such simulations give CCS values significantly larger than the experimental ones. Independent trajectories, started from different conformations and with different protonation states (*i.e.*, different sets of localized charges), confirmed this result (supporting Figure S7). The experimental CCS values of the duplexes^6-^ are nonetheless compatible with the simulated A- and B-helices, and duplexes^7-^ are compatible with B-helices. This does not necessarily mean that the gas-phase conformations of 6- and 7- charge states *are* A- and B-helices, but helps to understand conclusions previously derived solely based on more densely charged duplexes.^17,18^

When starting from the B-form, MD simulations always show spontaneous hydrogen bonds formation between phosphate groups situated on each side of the minor groove (supporting information Movie S1 and Figure S8). This causes the “zipping” of the minor groove. The structures are stable up to 1 μs (Figure S8), but too elongated compared to the duplexes^5-^ experimental data (compare Fig. 1B with 1A). In simulations starting from the A-helix, zipping occurs as well, but this time across the major groove (supporting Figure S9). In the gas phase, the closest protonated phosphate groups therefore tend to form hydrogen bonds that did not exist in solution. However, this kind of trajectory does not reflect an experimentally relevant one.

To check for possible artifacts due to using classical force fields to represent macromolecules in the gas phase, we also studied B-duplexes at the density functional theory (DFT) and semi-empirical (SE) levels. Upon DFT optimization of 7-bp to 10-bp duplexes, phosphate groups form new H-bonds and close the minor groove as well (Figure 2 and Supporting Figure S11). *WC* H-bonds and base stacking are well preserved,^37,38^ and the helix compresses along its longitudinal axis (Figure 2C) and the ^CALC^CCS_He_ of the DFT optimized structure is smaller than that of the canonical helix.

**Figure 2.**
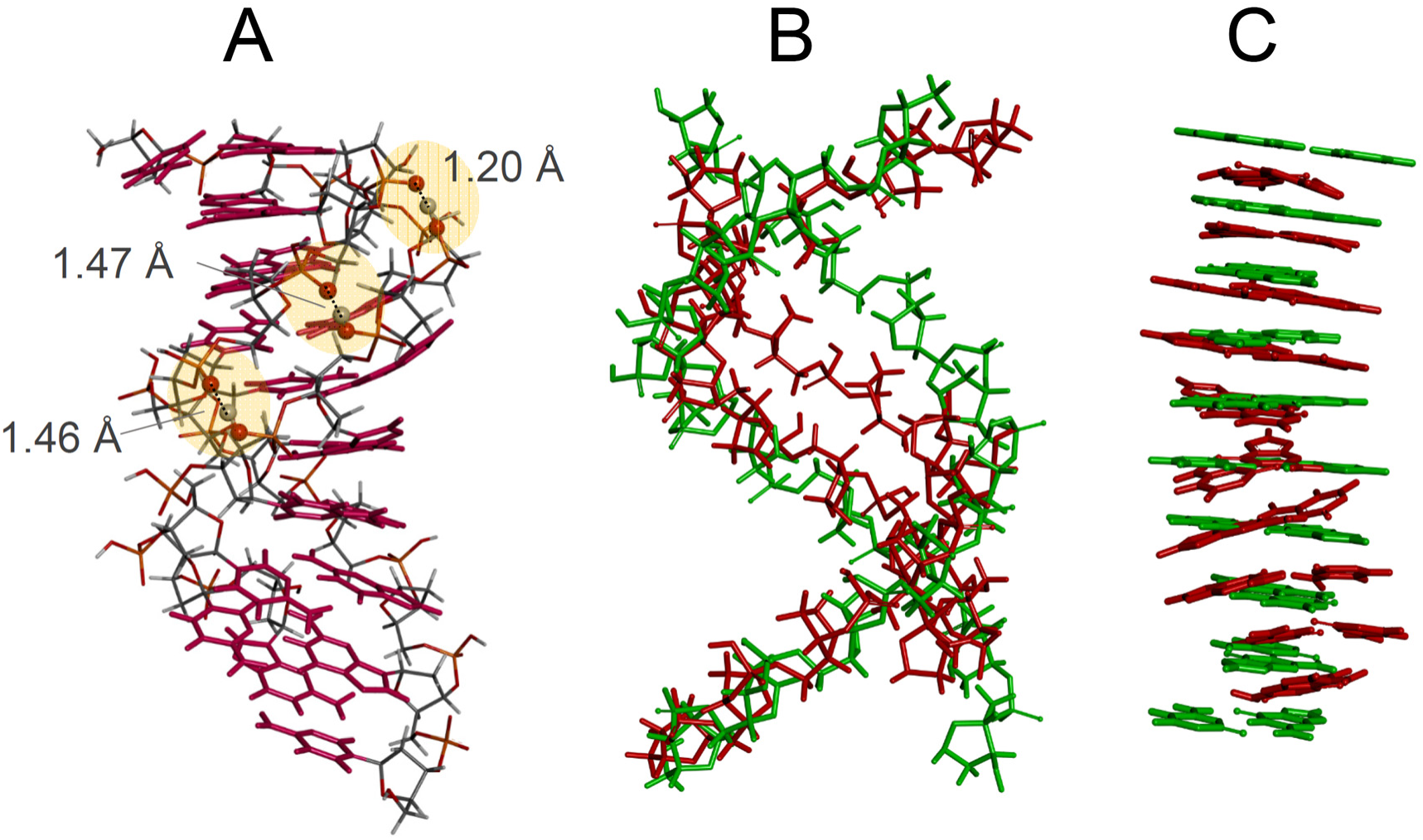
DFT optimization of the 10-bp duplex^4-^: (A) new hydrogen bonds formed between phosphate groups across the minor groove; (B) superposition of the sugar-phosphate backbones of the canonical B-helix (green) and optimized duplex (red); (C) superposition of the base pairs of the canonical B-helix (green) and optimized duplex (red).

For SE calculations, we first validated that the PM7 method best reproduces the DFT results (supporting Figure S12). The CCS values obtained after PM7 optimization are summarized in Table 2 for all [12-bp duplexes]^5-^ (A- and B-form for DNA and A-form for RNA). Upon PM7 optimization, the duplex [12-d_66_]^5-^ undergoes minor groove zipping, while *WC* H-bonds and base pair stacking interactions are preserved (Figure S13). The compaction compared to the solution structure is only ~8%, thus still far from the experimental value.

Thus, starting from naked canonical structures, neither geometry optimization nor unbiased MD trajectories lead the system towards the experimentally observed conformational ensemble. So, either the sampling is incomplete (simulated and experimental time scales differ), or the starting structures, obtained by desolvating and charging the duplexes all at once, inadequately reflect the electrospray droplet desolvation and declustering.

### 3) Temperature replica exchange MD (T-REMD) exploration of gas-phase conformational landscapes do not converge to the experimental structure.

To enhance sampling, we carried out T-REMD simulations on the 12-bp duplexes^5-^ starting from their canonical structures. We used both DC and LC models. DNA and RNA results, including representative snapshots, total hydrogen bonding, *WC* hydrogen bonding and stacking^39^ occupancies, are summarized in supporting information Section S5 (Figures S14—S24). For RNA, the theoretical CCS distributions, both with the DC and LC models, match with the experimental ones (Figures S15 and S18). However, the hydrogen bonding and stacking patterns (Figures S19-S21) always become scrambled in the gas phase. For DNA, the T-REMD results depend more critically on the charge location model: ^CALC^CCS_He_ values agree with the experiments for the DC model (Figure 1D) and for *some* LC models (Figure 3).

**Figure 3.**
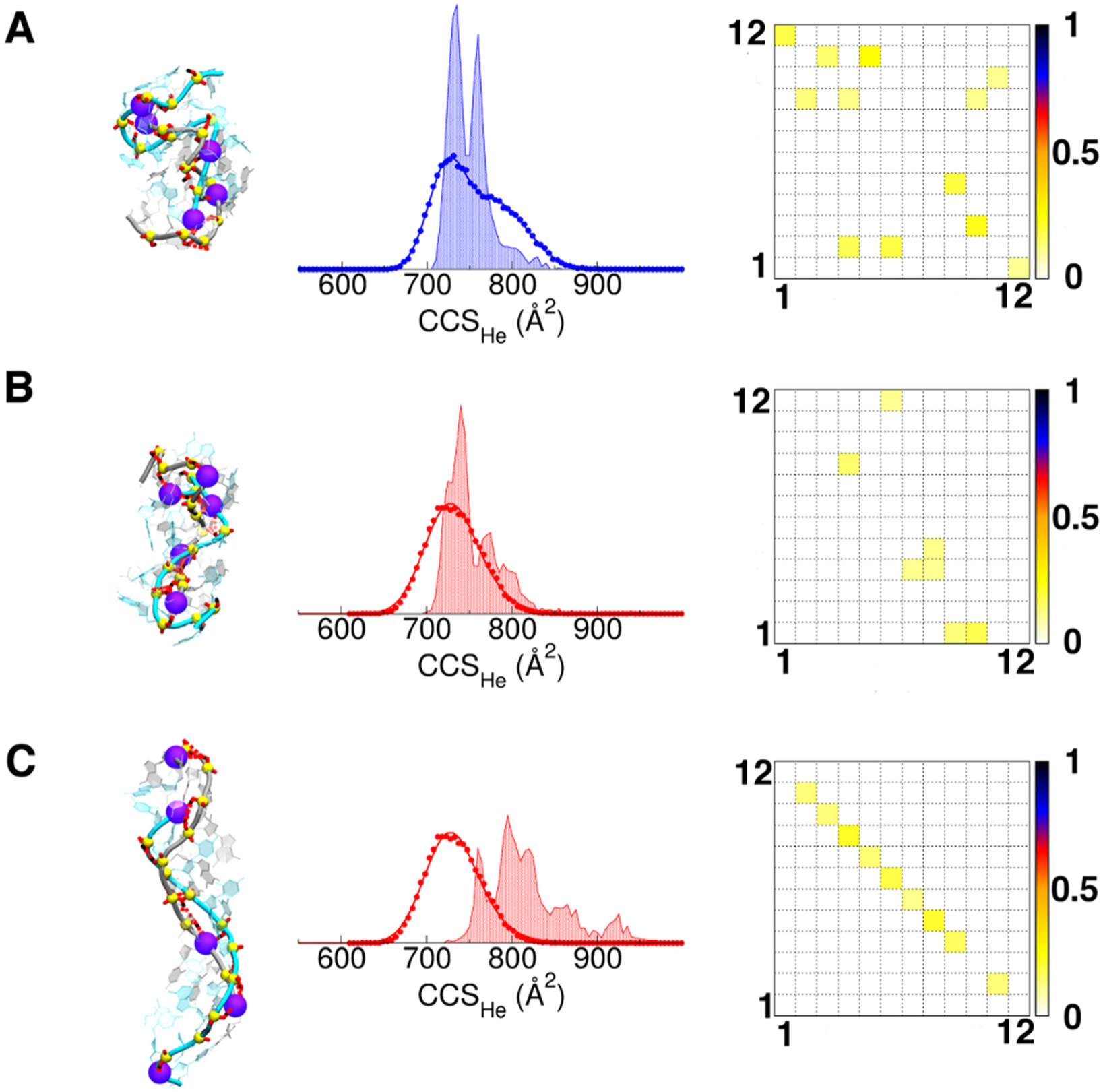
T-REMD on 12-bp DNA duplexes^5-^ with localized charges (LC). (A) [12-d_66_]^5-^, (B-C) [12-d_100_]^5-^ with different LC models. The location of negatively charged phosphates is shown by the beads on the representative snapshot structure. The experimental ^DT^CCS_He_ distribution and calculated one (shade) are overlaid. The fractional *WC* H-bond occupancies are shown on the right: the bases of each strand are numbered from 5’ to 3’, and perfect base pair matching is therefore indicated by a diagonal from top left to bottom right.

With the DC model, the CCS values are closer to the experimental ones, yet still significantly too large for 12-ds_100_^5-^. The GC base pairs are mostly preserved (Figure S36), but the duplex 12-d_33_^5-^ is mostly melted (separated into single strands; see Figure S14). DC models lack the explicit protons and therefore cannot form phosphate-phosphate hydrogen bonds. As a result, the strands can dissociate upon T-REMD. The rate of strand dissociation occurrence ranks 12-d_33_ > 12-d66 > 12-d_100_ (supporting Figure S14), in line with the relative gas-phase kinetic stabilities in tandem mass spectrometry.^11-14^

When LC models are used, the T-REMD final structures depend significantly on the choice of charge location (Figure S17, S19—S21). For example, for [12-d100]^5-^, a first model (Figure 3B) gives CCS values matching well with the experiment, but has lost most *WC* H-bonds, whereas a second model (Figure 3C) preserves *WC* H-bonds but its CCS values are much larger than the experimental ones (the representative structure resembles those obtained with unbiased simulations starting from the B-form).

In summary, the T-REMD results do not account simultaneously for the preservation of GC base pairs and the experimental collision cross sections. Yet they teach us that the duplex^5-^ conformations are closer to a compact globular shape than to the helices obtained by unbiased MD trajectories, and suggest that sampling problems are at least partially responsible of the unfitting of simulation and experimental CCSs.

### 4) Progressive duplex desolvation leads to experimentally relevant structures.

All simulations presented to this point, and nearly all previous simulations are assuming an instantaneous transfer of the duplex from solution to the gas phase. But in practice, desolvation and declustering proceeds gradually during the experiment. Consta and co-workers^40^ modeled the desolvation process of duplex dA_11_•dT_11_ at atomistic level in a water droplet containing Na^+^ and Cl^-^, and found that the duplex collapses inside the droplet when the Na^+^ cations are numerous enough to interact with the phosphates and reduce the size of both grooves. Interestingly, the resulting charge density is similar to those observed experimentally from native conditions. It is therefore likely that the DNA duplexes are desolvated via the charged residue model^41,42^ and that the compaction results from the association of NH_4_^+^ cations to both minor and major groove before full desolvation. As a result, the starting structure for gas phase simulations might be quite different to the canonical A- or B- helices. Because electrostatic interactions prevail, the gas phase conformational space is very stiff, and an incorrect starting structure can significantly bias the entire trajectory.

Here we simulated the desolvation of duplex 12-d_66_ placed in a droplet containing ~2400 water molecules and 17 NH_4_^+^ (net charge = −5). When the simulation reaches 39.5 ns, 87 water molecules stick on the duplex together with the 17 NH_4_^+^ cations. The simulation was pursued for 50 ns at 450 K, to allow this ultra-stable inner solvation shell of water molecules to evaporate. The ammonium ions sticking to the duplex are mainly located close to phosphate groups, often in-between them. Several trajectories (see 4-, 5- and 6- charge states in supporting information Section S6; Figures S25—S30) lead to a chain of ammonium ions in the minor groove. For the 5- charge state, ammonium ions are more distributed across both grooves, and thus upon evaporation both grooves get narrower to enable phosphate-ammonium-phosphate salt bridges to form. Accordingly, the CCS value diminishes (Figure 4).

**Figure 4.**
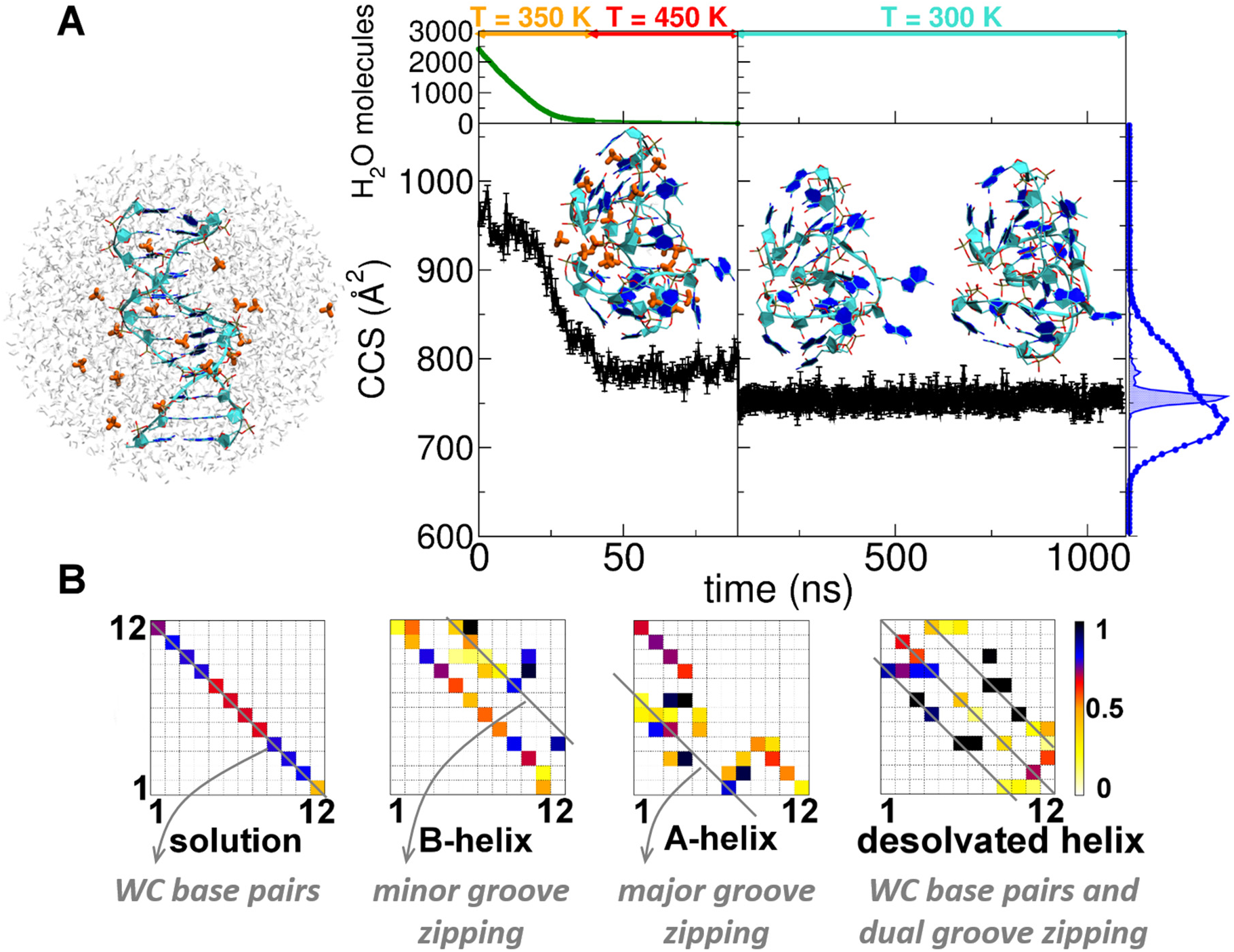
(A) ^CALC^CCS_He_ evolution during the desolvation of the 12-d66 droplet with −5 net charge. At the top the simulation temperatures are shown along the related trajectory portions. The initial, intermediate and final structures are shown with DNA strands in cyan, bases aromatic rings in blue and NH_4_^+^ cations in orange. On the right the [12-d66]^5-DT^CCS_He_ distribution (blue circles) is superimposed on the CALCCCS_He_ (blue area) one. (B) H-bond occupancies of the desolvated helix (right), compared to Watson-Crick H-bond occupancies of a B-helix in solution (left), and to gas-phase B- or A- helices upon unbiased MD in the gas phase (middle). In the B-helix, extra H-bonds form between phosphates across the minor groove; in the A-helix, across the major groove, and in the desolvated helix, across both grooves.

Classical MD cannot model proton transfer, but if water evaporation is almost complete before the proton transfers start, then the structures generated by gradual desolvation are good candidate for modeling the electrosprayed structures. Also, ammonium ion positioning upon desolvation could predict which phosphate groups will share a proton and form hydrogen bonds after complete declustering.

To simulate the eventual ammonia loss, we arbitrarily transferred a proton from each ammonium ion to its closest phosphate oxygen. The resulting desolvated and declustered duplex (12-d_66_)^5-^ is stable over 1-µs MD at 300 K (Figure 4). The total hydrogen bond occupancy reveals additional contacts across both grooves. Remarkably, the CCS value now matches the experimental one, and at the same time, the generated structure keeps partial memory of the *WC* base pairs (at least, preserving mostly the GC ones), in line with CID and IRMPD results.

In summary, progressive desolvation, allowing the duplex to form phosphate-phosphate hydrogen bonds across both grooves can account simultaneously for the experimental compactness and for partial preservation of GC base pairs. In contrast, upon unbiased MD or structure optimization (DFT or SE), only the phosphate groups that were the closest in the starting structure (across the minor and major groove for B- and A-helices, respectively) could mate.

### 5) Longer duplexes, DNA and RNA alike, undergo compaction when electrosprayed from aqueous solutions of physiological ionic strength.

Past studies on GC-rich DNA duplexes, with one charge every two base pairs, had suggested that A-helices predominate from 8-bp on, and that B-helices are preserved from 18-bp on. To ascertain whether the compaction of lower charge states is a general phenomenon, we measured 12-bp to 36-bp DNA and RNA duplexes, from native solutions, either 100% aqueous or containing sulfolane. The tested duplexes included multiples of d_66_, r_66_, d_100_ and r_100_, and the DNA duplexes listed as potential CCS calibrants by the Fabris group.^43^ The results are shown in Figure 5 (experimental values in supporting Table S3).

**Figure 5.**
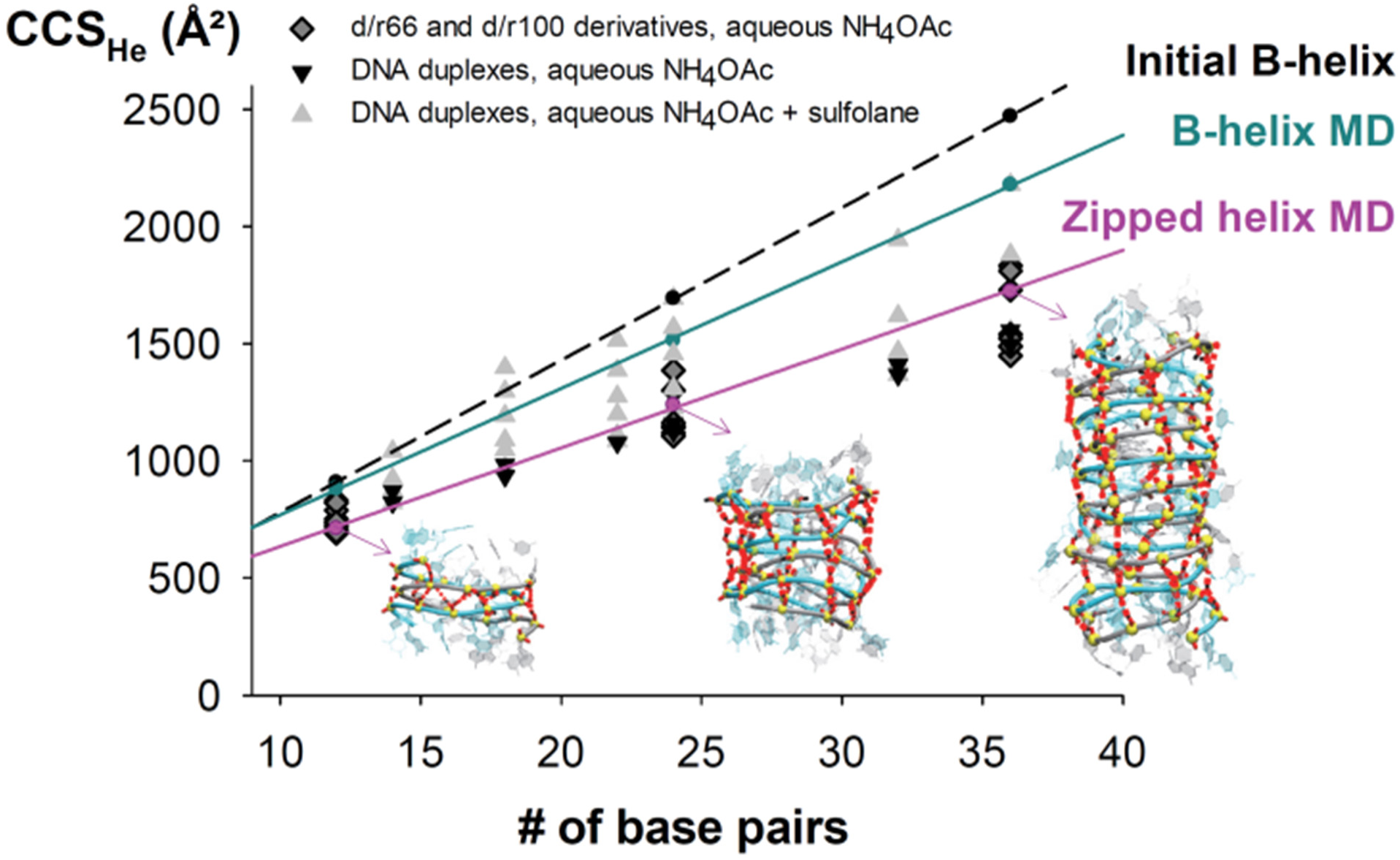
Comparison between experimental and calculated CCS_He_ of 12-bp to 36-bp duplexes. Diamonds and black down triangles: experiments on DNA and RNA duplexes in aqueous 100-mM NH_4_OAc, respectively. These experimental points match better with MD on the zipped helix (restrained minimization followed by MD). DNA duplexes at higher charge states produced by sulfolane addition adopt more extended conformations, which match better with MD simulations on B-helices.

We found that DNA and RNA duplexes have very similar gas-phase CCS values, although their helix types differ in solution (B *vs.* A). The theoretical CCS values obtained with unbiased *in-vacuo* MD directly from the solution B-helix are overlaid (B-helix MD trend line in Fig. 5). They match only with some of the high charge states produced in the presence of sulfolane. However, the low charge states produced from purely aqueous NH_4_OAc solutions are significantly more compact.

To reproduce the dual-groove closing of longer duplexes while avoiding computationally costly desolvation simulations, we opted for biased exploration. We used restrained minimization and imposed distance constraints based on our knowledge that compact structures can form via hydrogen-bond formation between phosphates across both grooves. Distance restraints were imposed using a harmonic potential between the hydrogen atom of neutral phosphates belonging to one strand and the oxygen atom of mating phosphates belonging to the other strand (Supporting Information Section S8, Figures S31—S33, and Movie S2 for a 12-bp duplex). The systems were minimized *in vacuo*, then all restraints were removed prior to 1-μs gas phase MD.

For 12-bp duplexes, the resulting CCS distributions agree with the experiment, with the strongest compaction for 12-d_100_ (Figure 1E). Applying the same procedure starting from an A-helix however did not lead to similarly low CCS values (supporting Figure S34). The doubly groove-zipped helices obtained by restrained minimization generally keep *WC* hydrogen bonds less well than those obtained by desolvation (supporting Figure S35—S37), but still reflect the solution trend, with the 12-d_100_ preserving the highest fraction of hydrogen bonds. The advantage of the procedure is to reproduce the phosphate-phosphate H-bond pattern of the desolvated helices (two diagonals for zipping across both grooves, plus the central diagonal indicating preserved base pairs, Figure S35). We then applied restrained minimization to the longer (24-bp and 36-bp) helices (supporting Figures S38—S39). Whatever the duplex length, the experimental CCS values obtained for low charge states (Figure 5) match better with these zipped helices than with the canonical structures or with the helices relaxed by long unbiased gas-phase MD.

## CONCLUSIONS

In summary, at the charge states produced by electrospray from aqueous ammonium acetate (traditional “native” solution conditions), double-stranded nucleic acids undergo a significant compaction in gas phase compared to the structure in solution. Unbiased molecular dynamics of B-helix or A-helix structures directly transposed from solution to the gas phase fails to reproduce the experimental results. This is due to several reasons: i) only the phosphate groups closest to one another can pair to form hydrogen bonds on the simulation time scale, ii) the starting structure is unrealistic and iii) sampling in unbiased MD simulations is intrinsically limited. In the case of T-REMD, depending on the initial choice of charge location, the final structures either did not have any memory of the solution structure, or resembled those obtained by unbiased MD. TREMD can help to solve the sampling effect (the question of the maximum internal temperature reached in the experiments remaining open), but not the problem that original charge locations might be incorrect.

Gradual desolvation generates more realistic starting structures for gas phase simulations. Conformational transitions occurring during dehydration cannot be ignored because they guide the entire sampling, within a particularly stiff conformational landscape in the case of nucleic acids. The conformationally restrained duplexes remain stable upon unbiased MD: once formed, they stay locked at room temperature. The broadness of the experimental CCS distributions therefore indicates a distribution of co-existing—but not interconverting—conformations, wherein each would have a slightly different phosphate-phosphate hydrogen bond network.

Our results highlight a key difference between nucleic acids and proteins native mass spectrometry. Globular proteins can rearrange by relaxing their side chains^44^ and undergo minimal salt bridge rearrangement.^45^ Briefly optimized structures often have CCS values matching well with the experiments.^42,46-48^ Fabris and co-workers have recently underlined the difficulties in transposing to DNA the MD and CCS calculation protocols traditionally used for proteins.^43^ As a way out they proposed to calibrate all traveling wave IMS data using short MD simulation results, but our study shows why this approach would lead to a misrepresentation of nucleic acid structures in the gas phase. DNA and RNA double helices are more compact in the gas phase than in solution, due mostly to new phosphate-phosphate interactions. At the low charge states produced from ammonium acetate, the Coulomb repulsion is not sufficient to keep the phosphate groups apart. They rearrange by self-solvation, cause major rearrangements of the backbone, and lead to a significant compaction (>20%) compared to the starting structure. Yet, they are metastable conformations keeping some memory of what the structure was just before vaporization.

## SUPPORTING INFORMATION

Experimental procedures including reconstruction of the CCS distributions, full ESI-IMS-MS results, detailed computational procedures, full modeling results by *in vacuo* QM optimization, MD, T-REMD, and MD following restrained minimization, structural analysis, and supplementary results on longer duplexes (PDF).

Movie S1: minor groove zipping upon MD of a 12-bp B-helix (GIF)

Movie S2: dual groove zipping imposed by restrained minimization on a 12-bp B-helix (GIF).

## ACKNOWLEDGMENTS

This work was funded by the European Research Council (ERC DNAFOLDIMS to VG and ERC SimDNA to MO), by the Spanish BIO2015-69802-R grant to MO, and by EU COST action BM1403 (short-term scientific mission to MP). HG is a Juan de la Cierva researcher.

We are thankful to David E. Condon for donating us the Perl script used to score the base stacking, and to Adrien Marchand and members of COST action BM1403 for fruitful discussions. LD is a SNI (Sistema Nacional de Investigadores; ANII, Uruguay) researcher, and MO is an ICREA-academia fellow.

## ABBREVIATIONS

bp: base pair
CCS: collision cross section
^CALC^CCS_He_: calculated CCS in helium
^DT^CCS_He_: CCS in helium measured in a drift tube
DC: distributed charges
DFT: density functional theory
ESI: electrospray ionization
GC: guanine-cytosine
IMS: ion mobility spectrometry
LC: localized charges
MD: molecular dynamics
MS: mass spectrometry
SE: semi-empirical
TREMD: temperature replica exchange molecular dynamics
*WC*: Watson-Crick.

